# An In situ Collagen-HA Hydrogel System Promotes Survival and Preserves the Proangiogenic Secretion of hiPSC-derived Vascular Smooth Muscle Cells

**DOI:** 10.1101/2020.06.06.137968

**Authors:** Biraja C. Dash, Kaiti Duan, Hao Xing, Themis R. Kyriakides, Henry C. Hsia

## Abstract

Human induced pluripotent stem cell-derived vascular smooth muscle cells (hiPSC-VSMCs) with proangiogenic properties have huge therapeutic potential. While hiPSC-VSMCs have already been utilized for wound healing using a biomimetic collagen scaffold, an in situ forming hydrogel mimicking the native environment of skin offers the promise of hiPSC-VSMC mediated repair and regeneration. Herein, the impact of a collagen type-I-hyaluronic acid (HA) in situ hydrogel cross-linked using a PEG-based cross-linker on hiPSC-VSMCs viability and proangiogenic paracrine secretion was investigated. Our study demonstrated increases in cell viability, maintenance of phenotype and proangiogenic growth factor secretion, and proangiogenic activity in response to the conditioned medium. The optimally cross-linked and functionalized collagen type-I/HA hydrogel system developed in this study shows promise as an in situ hiPSC-VSMC carrier system for wound regeneration.

## Introduction

Human-induced pluripotent stem cells (hiPSCs) have shown enormous potential for applications in regenerative therapy and disease modeling (Shi, Inoue, Wu, & Yamanaka, 2017; Soldner & Jaenisch, 2012; Takagi et al., 2019; Takahashi & Yamanaka, 2006). Human iPSCs with properties similar to embryonic stem cells can be differentiated into various specialized cells such as endothelial (hiPSC-ECs) and vascular smooth muscle cells (hiPSC-VSMCs) (Ayoubi, Sheikh, & Eskildsen, 2017; B. C. Dash, Z. X. Jiang, C. Suh, & Y. B. Qyang, 2015b; Klein, 2018; Maguire, Xiao, & Xu, 2017). The hiPSC-VSMCs in pure population have been generated in abundance (Cheung, Bernardo, Trotter, Pedersen, & Sinha, 2012; Dash et al., 2015b; Dash et al., 2016; Patsch et al., 2015) and their ability to carry the disease mutations and to secrete collagen has been used to develop robust disease models (Atchison et al., 2020; Atchison, Zhang, Cao, & Truskey, 2017; Dash et al., 2016; Ge et al., 2012; Liu et al., 2011; Zhang et al., 2011) and vascular grafts (Gui et al., 2016; Karamariti et al., 2013; Luo et al., 2020), respectively. In addition, we recently reported the secretion of various growth factors and cytokines from these cells (Dash et al., 2020; Gorecka et al., 2020). Specifically, hiPSC-VSMCs, when embedded in a dense collagen scaffold, experienced a microenvironment attributable to a combination of mechanical forces and hypoxia that induced secretion of growth factors and cytokines including vascular endothelial growth factor (VEGF), basic fibroblast growth factor (bFGF), angiopoietin-1 (ANG-1), transforming growth factor (TGF)-β, platelet-derived growth factor (PDGF), matrix metalloproteinase (MMP)-2, stromal cell-derived factor (SDF)-1α, interleukin (IL)-10 and IL-8 (Dash et al., 2020). Moreover, hiPSC-VSMCs in vivo exhibited proangiogenic and anti-inflammatory activities and were shown to promote regenerative healing in both acute and diabetic wound models in nude mice (Dash et al., 2020; Gorecka et al., 2020). These initial studies have shown future translation potentials of hiPSC-VSMCs in wound healing and ischemic diseases.

Extracellular matrix (ECM)-based biomaterials such as collagen have been designed for the delivery of stem cells’ beneficial properties (Dash et al., 2018; Hinderer, Layland, & Schenke-Layland, 2016). While collagen and its biophysical properties have been shown to have significant impact on stem cells’ therapeutic function, cross-linked in situ collagen-based hydrogels have been developed to enhance their stability and ease of application (Ahmed & Ffrench-Constant, 2016; Dash et al., 2020; Hou, Coller, Natu, Hastie, & Huang, 2016; Watt & Huck, 2013). Widely used chemical cross-linking methods have impact on collagen matrix architecture, stability, porosity, morphology, and function of stem cells (Depalle, Qin, Shefelbine, & Buehler, 2015; Gu, Shan, Ma, Tay, & Niu, 2019). Polyethylene glycol (PEG)-based cross-linker has been used to develop mechanically robust and enzyme resistant biomaterials without compromising their biocompatibility (Lee, Tong, & Yang, 2016; Taguchi et al., 2005). Previously, 4arm PEG Succinimidyl Glutarate (4S-StarPEG) has been used to crosslink collagen type-I and II to fabricate various forms of cell and drug delivery systems (Browne et al., 2015; Collin et al., 2011; Fontana et al., 2015; Fontana, Thomas, Collin, & Pandit, 2014; Monaghan, Browne, Schenke-Layland, & Pandit, 2014; Thomas et al., 2014). 4S-StarPEG-based carrier systems with enhanced mechanical strength have been used to provide a niche for survival and regenerative function of mesenchymal stem cells (MSCs), adipose-derived stem cells (ADSCs), nucleus pulposus cells (NPCs) (Collin et al., 2011; Fontana et al., 2015; Fontana et al., 2014; Thomas et al., 2014).

Hyaluronic acid (HA), a non-sulfated glycosaminoglycan, is abundantly present in the ECM of the skin and many other major organs and has tissue-specific functions (Papakonstantinou, Roth, & Karakiulakis, 2012). HA has been shown to promote cellular proliferation and migration, and blood vessel formation during tissue injury (Gerecht et al., 2007; Hanjaya-Putra et al., 2012; Kwon et al., 2019; Shen et al., 2014). HA contains cell-binding motifs and has been widely used for surface modification of biomaterials to support cell survival and proliferation (Highley, Prestwich, & Burdick, 2016; Khetan & Corey, 2019; Solis et al., 2012). Several earlier studies have reported the use of HA-based hydrogels to maintain stem cells such as NPCs, ADSCs, and hiPSC-derived vascular cells (Collin et al., 2011; Fontana et al., 2014; Shen et al., 2016). In particular, one of the studies that used acrylated HA-based hydrogel for hiPSC-derived early vascular cells, demonstrated enhanced angiogenic potentials that translated to improved wound healing capacities in diabetic mice (Shen et al., 2016).

In this study, we characterized an in situ cell-carrier system for hiPSC-VSMCs. The biomaterial system was comprised of type I collagen hydrogel optimally stabilized with 4S-StarPEG and enriched with HA. In addition, we investigated how collagen density is a crucial factor impacting chemical and biophysical behavior of the in situ hydrogels, and ultimately cellular viability, paracrine function, and proangiogenic potentials. It is our hypothesis that a system mimicking skin composition would be a suitable carrier for hiPSC-VSMCs for wound healing. Specifically, an in situ hydrogel will allow encapsulation and delivery of hiPSC-VSMCs, and provide a better conducive environment for the maintenance of their cell viability, paracrine functions, and proangiogenic therapy.

## Methods

### In situ collagen-HA hydrogel fabrication

The in situ hydrogels were fabricated using a rat tail collagen type-I, HA and 4S-StarPEG (Schematic 1). Briefly, the in situ collagen-HA hydrogels were generated by properly mixing collagen type-I of initial concentration of 5mg/ml, 10xMEM, and 1M NaOH followed by the addition of final concentration of 1mg/ml of HA and cross-linker 4S-StarPEG. Various molar ratios (1:0.5, 1:1 and 1:2) of collagen to the 4S-StarPEG cross-linker were tested. The entire hydrogel generation process was done on ice. The mixture ratios of varying collagen densities to 10x modified essential medium (MEM), 1M NaOH, HA, and 4S-StarPEG has been listed in the tables in the supplementary information. The resultant mixture was mixed quickly and gently to avoid bubble formation and quick gelation. The collagen mixture was then distributed as 100µL aliquots into 96 well cell culture plate and incubated for 30 minutes at 37°C for the gelation. The final in situ collagen-HA hydrogel contained 4mg/ml and 1mg/ml final concentration of collagen and HA respectively and 1:1 molar ratio collagen to 4S-StarPEG cross-linker. The controls were cross-linked collagen hydrogels and uncross-linked collagen and collagen-HA hydrogels. Please refer to the tables (Table S1-S3) in the supplementary information for a detailed composition of each of these hydrogels.

### Cell seeding

HUVECs and dermal fibroblasts: HUVECs and dermal fibroblasts from primary sources were seeded on top of the hydrogels at a concentration of 1×10^5^/ml of cells. The primary cells were cultured for 3 days supplemented with 100µl of the respective medium on the top of the hydrogels. The hydrogels were tested on days 1 and 3 for cell viability.

hiPSC-VSMCs: hiPSC-VSMCs of 4 × 10^5^/ml concentration were embedded while forming the collagen-HA in situ hydrogels. The cells were added after HA and before the cross-linker during the process. The hiPSC-VSMCs were cultured for 3 days supplemented with 200µl of SmGM-2 medium on the top of the hydrogels. After day 3, the hydrogels were tested for cell viability and phenotype while the 200µl of conditioned medium (CM) was collected to analyze paracrine secretion and proangiogenic function.

## Results

### Cross-linking and cell viability of in situ collagen hydrogel is collagen density dependent

4S-StarPEG was used to fabricate in situ collagen hydrogels as reported earlier. We used rat tail Type-I collagen of 5mg/ml initial concentration to develop hydrogels of the final concentration of 1.25, 2.5 and 4mg/ml. TNBSA assay was used to evaluate cross-linking by determining the amine content of the hydrogels (Figure 1). The initial amount of amine was determined for un-crosslinked hydrogels to determine the collagen to 4S-StarPEG ratios of 1:0.5, 1:1 and 1:2. TNBSA data showed a decrease in the free amine groups in the cases of cross-linked hydrogels and was dependent on their final collagen concentration. For 1.25mg/ml of hydrogels only 1:2 ratio had a significant reduction in the level of the amine group and was reduced compared to control 1:0.5 and 1:1, while no difference was seen between control, 1:0.5 and 1:1 (Figure 1A). However, in the case of 2.5mg/ml 1:2 there was higher degree of cross-linking than 1:0.5 and control groups, and 1:1 ratio showed higher cross-linking compared to the control group. There were no significant differences between 1:2 and 1:1, 1:1 and 1:0.5, and 1.05 and control (Figure 1B). For the 4mg/ml group 1:2, 1:1 and 1:05 were all more cross-linked than control, and 1:2 was higher than 1:0.5. But there were no differences between 1:1 and 1:0.5, and 1.2 and 1:1 (Figure 1C). The percentage of cross-linking was found to be higher in the case of 4mg/ml hydrogels with a 40-70% reduced amine group compared to 20-50% in 2.5mg/ml and 7-20% in 1.25mg/ml hydrogels (Figure 1D). Collagen cross-linked with glutaraldehyde was used as a positive control and TNBSA assay showed reduced free amines (Figure S1).

**Figure 1:**
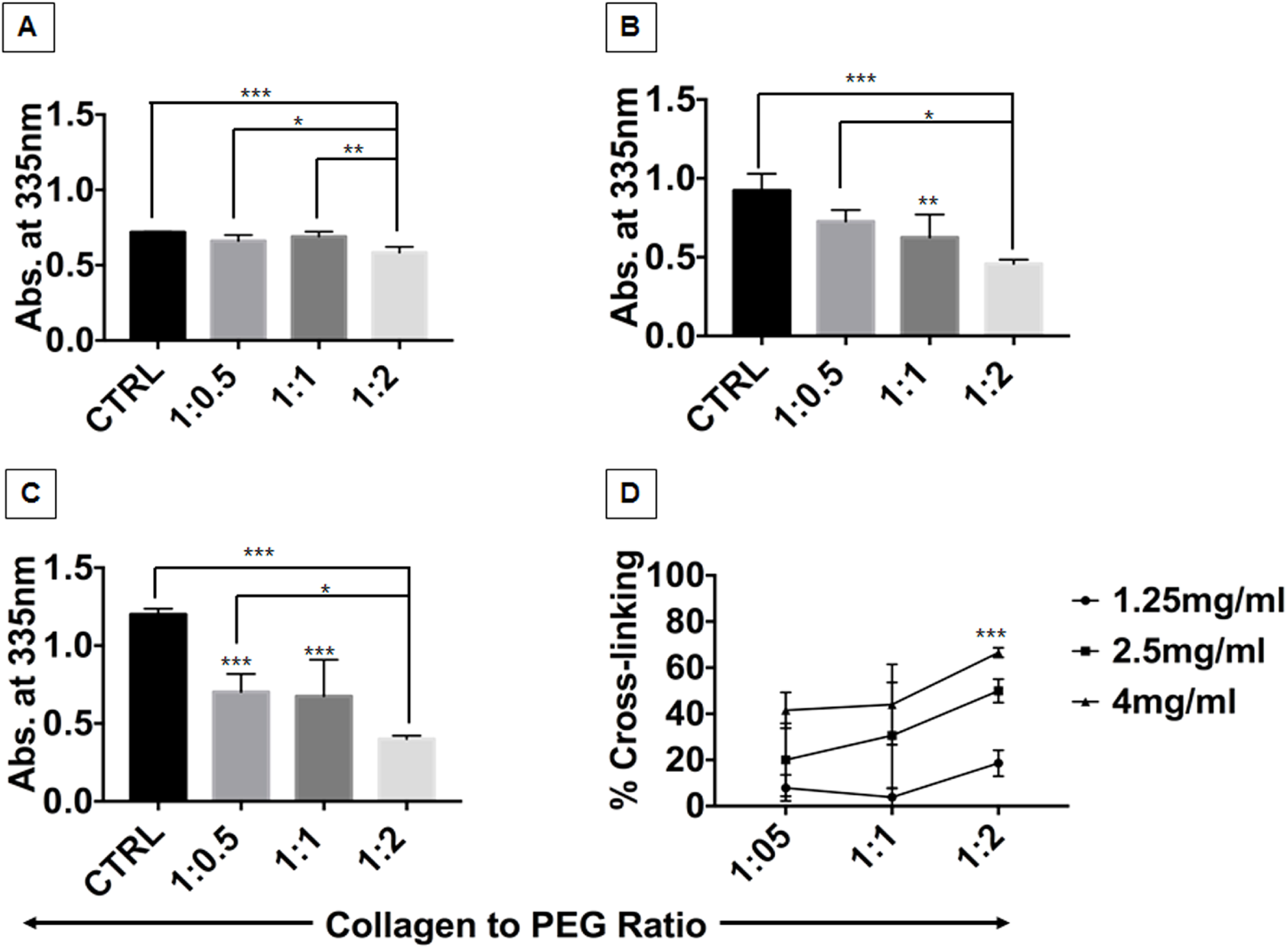
Characterization of 4S-starPEG cross-linking of collagen type-I hydrogels. TNBSA showing qualitative analysis of free amine groups in in situ collagen hydrogels of (A) 1.25mg/ml, (B) 2.5mg/ml and (C) 4mg/ml of collagen concentration with various molar ratios (1:0.5, 1:1, and 1:2) of collagen to 4S-starPEG. (D) The cumulative data presenting percent cross-linking of various hydrogels. * denotes statistical significance differences between the different groups (n=4, one-way ANOVA, *p < 0.05, **p<0.005, ***p<0.001, and ****p<0.0001). Control (CTRL) is Uncross-linked collagen hydrogel.

The biocompatibility of the 4S-StarPEG cross-linked collagen hydrogels was determined using HUVECs and dermal fibroblast cells (Figure 2). AlamarBlue assay was performed to assess the relative cell viability of dermal fibroblasts and HUVECs cultured on the hydrogels. On day 1 an increase in the cell viability of fibroblasts was seen with increasing collagen concentration irrespective of cross-linker ratios except for 1:2 where 2.5mg/ml was similar to 1.25mg/ml (Figure 2A). On day 3, an overall increase in the cell viability of fibroblasts compared to day 1 was observed (Figure 2B). 4mg/ml was found to be significantly higher compared to 2.5mg/ml and 1.25mg/ml across different cross-linker ratios. On both days 1 and 3, there was no difference in the cell viability of fibroblasts in relation to cross-linker ratios except control 4mg/ml day 3 was found to be higher than the 4mg/ml cross-linked groups (Figure 2B). Similarly, Day 1 HUVECs showed an increase in cell viability in 4mg/ml hydrogel groups irrespective of cross-linking ratios compared to other concentrations. However, unlike fibroblasts, on day 1 there was no difference between 2.5mg/ml and 1.25mg/ml across all the cross-linking ratios. Most importantly, on day 1 HUVECs on 4S-StarPEG cross-linked collagen hydrogels displayed an increase in viability with increasing cross-linking ratios except 1:2 (Figure 2C). Like dermal fibroblasts, we saw an overall increase in viability of HUVECs on day 3 compared to day 1 and an increase in viability in 4mg/ml hydrogel compared to all other groups. (Figure 2D).

**Figure 2:**
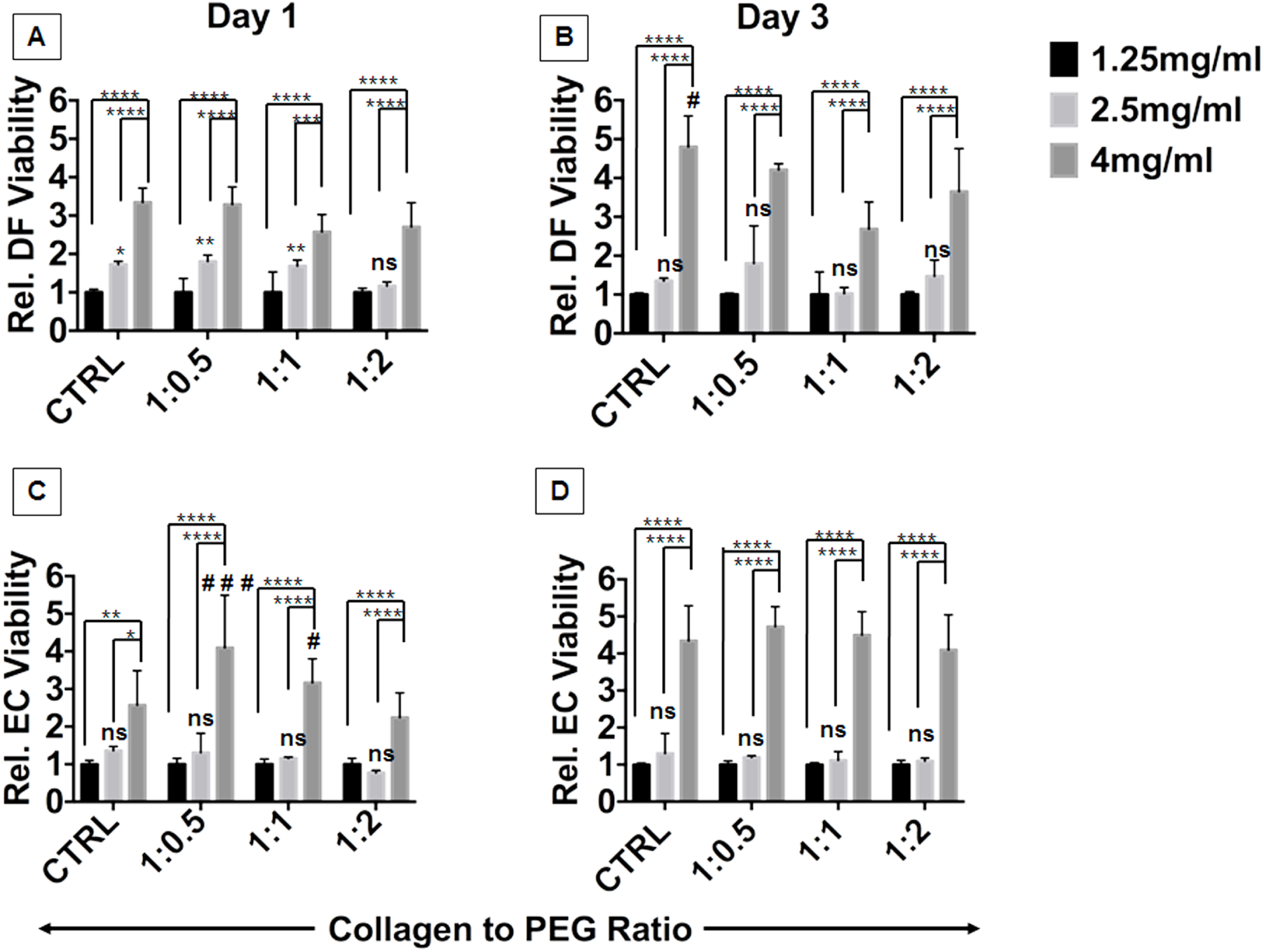
Characterization of biocompatibility of in situ collagen hydrogels. AlamarBlue assay showing relative cell viability of fibroblasts on (A) day 1 and (B) day 3 and HUVECs on (C) day 1 and (D) day 3 of culture on top of 1.25mg/ml, 2.5mg/ml and 4mg/ml of collagen concentration and different collagen to 4S-starPEG ratios. * and # denote statistical significance differences between the different groups (n=5, two-way ANOVA, *p<0.05, **p<0.005, ***p<0.001, ****p<0.0001, ^**#**^p<0.05, and ^**# # #**^p<0.001). Control (CTRL) is Uncross-linked collagen hydrogel.

hiPSC-VSMCs derived utilizing an EB method generated a pure population of synthetic VSMCs in abundance within 21 days. The hiPSC-VSMCs when characterized using synthetic VSMC markers calponin and SM-22α were found to be highly positive. These synthetic hiPSC-VSMCs were then embedded within the in situ hydrogels (Figure 3A). Cells embedded in different concentrations 4S-StarPEG of hydrogels were tested for viability using AlamarBlue. The data suggest no changes in the cell viability with increasing ratios of 4S-StarPEG in the case of 1.25mg/ml groups and 4mg/ml hydrogels (Figure 3B). However, in the case of 2.5mg/ml group, the viability of hiPSC-VSMCs increased with an increase in the cross-linking to 1:1 and 1:2., whereas 1:0.5 ratio was similar to control (Figure 3B). Overall, the cell viability was higher in the case of 4mg/ml hydrogels compared to 1.25 and 2.5mg/ml.

**Figure 3:**
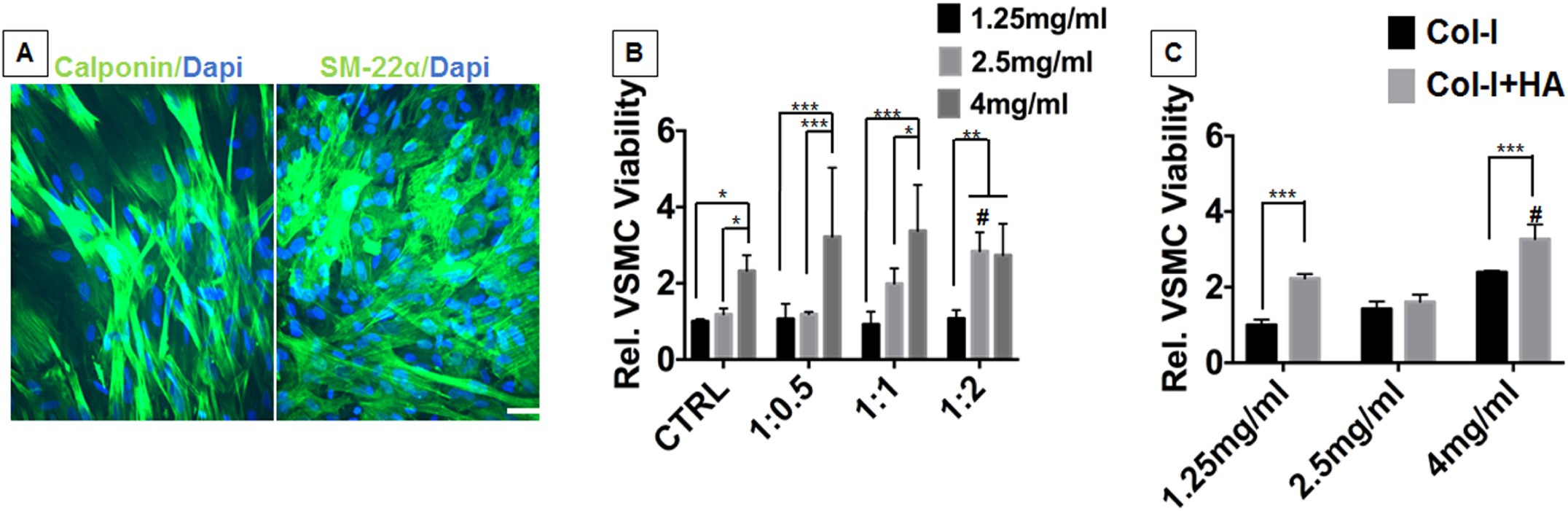
Characterization of the effect of cross-linking and HA on hiPSC-VSMC’s viability. (A) The hiPSC-VSMCs differentiated using an embryoid body method were characterized using smooth muscle cell markers. Phenotype assessment of hiPSC-VSMCs were performed using immunofluorescence staining with (A) Calponin (Green) and (B) SM-22α (Green). Dapi (Blue) was used to stain nuclei. Scale bar measures 43 µm. AlamarBlue assay showing relative cell viability of hiPSC-VSMCs embedded in the collagen hydrogels of 1.25mg/ml, 2.5mg/ml and 4mg/ml of collagen concentration with (B) various collagen to PEG ratios and (C) HA on day 3. * denotes statistical significance differences between the different groups (n=4, one-way ANOVA, *p < 0.05 and ****p < 0.0001).

### In situ Collagen-HA hydrogel promotes cell viability and maintains the paracrine function

HA of 1mg/ml final concentration was added while fabricating collagen hydrogels. The effect of HA on the viability of hiPSC-VSMCs embedded was evaluated using the alamarBlue assay. We observed an increase in the viability of hiPSC-VSMCs in 1.25mg/ml and 4mg/ml scaffolds (Figure 3C). However, we did not see any difference in the case of 2.5mg/ml group (Figure 3C). The difference in the increase in the viability was bigger in the case of 1.25mg/ml than 2.5mg/ml and 4mg/ml hydrogels (Figure 3C). 4mg/ml scaffold with 4S-StarPEG of 1:1 ratio and HA of 1mg/ml was chosen as the final hydrogel component (Figure 4). Gross images of the hydrogel showed an increase in transparency with 4S-StarPEG cross-linker (Figure 4A). This was further validated by quantifying % transmittance at 450 nm (Figure 4B). SEM was performed to analyze the surface morphology of hydrogels (Figure 4C). SEM images showed a fibrillar structure in the case of uncross-linked collagen hydrogels (Col-I) and the collagen-HA in situ hydrogels (Col-I+PEG+HA), whereas fibrillar structure was absent in the case of cross-linked collagen hydrogels (Col-I+PEG) (Figure 4C). The hiPSC-VSMCs embedded in the hydrogel showed a significant increase in cell proliferation with reduced cytotoxicity compared to Col-I and Col-I+PEG control hydrogels as shown in multiple cell viability experiments (Figure 5). AlamarBlue assay showed an enhanced cell proliferation in Col-I+PEG+HA hydrogels compared to Col-I hydrogel (Figure 5A), whereas LDH-based cytotoxicity assay showed a reduced level of LDH in both the CM and cell lysate in the case of both Col-I+PEG and Col-I+PEG+HA compared to Col-I uncross-linked hydrogels (Figure 5B and C). Live/Dead assay showed an increased number of live cells in Col-I+PEG+HA hydrogels compared to Col-I and Col-I+PEG control hydrogels (Figure 5D and E). The ratios of the dead to the total number of cells were found to be similar in all the hydrogels (Figure 5F).

**Figure 4:**
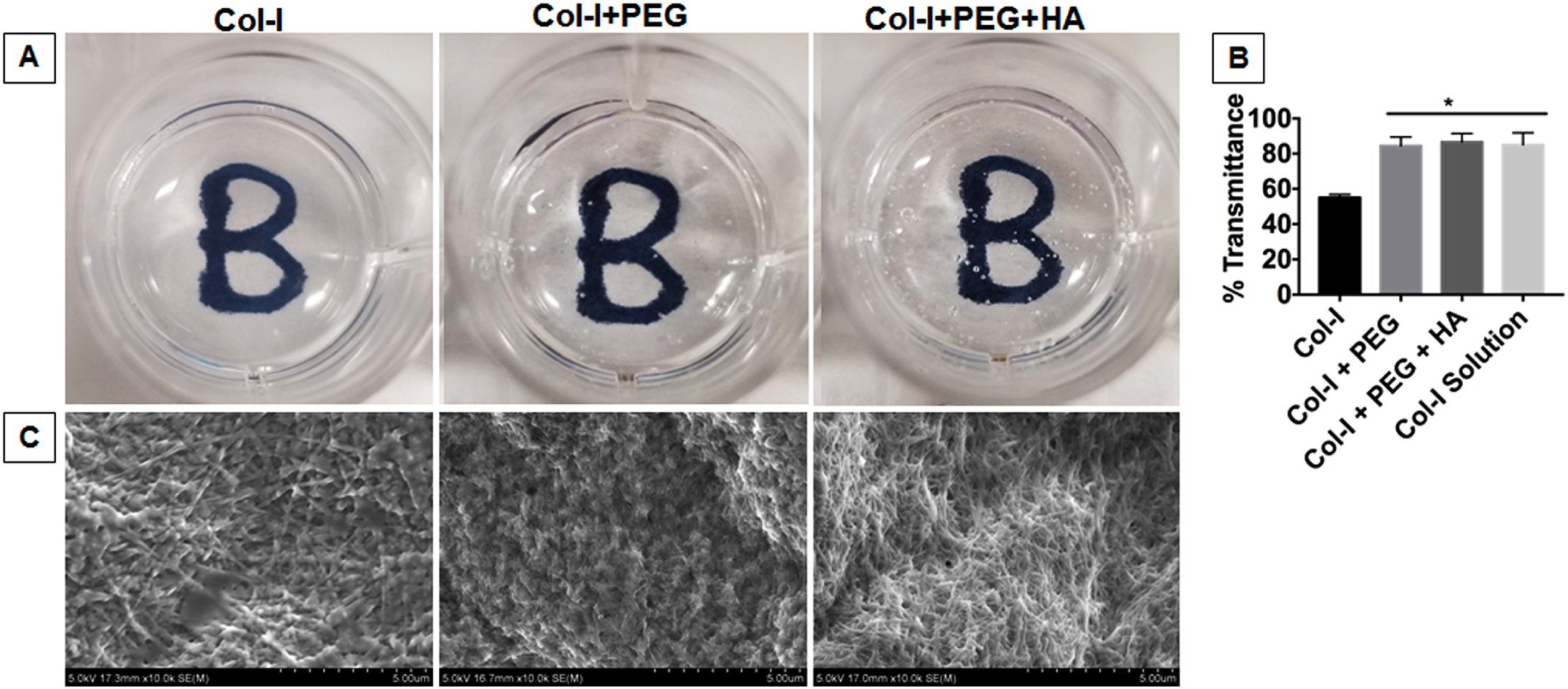
Biophysical characterization of in situ hydrogel. (A) Macroscopic images of uncross-linked collagen hydrogels (Col-I) and cross-linked collagen hydrogels with 1:1 ratio of collagen to 4S-StarPEG (Col-I+PEG) and cross-linked hydrogel with HA (Col-I+PEG+HA). (B) Transmission of Col-I, Col-I+PEG, Col-I+PEG+HA and Col-I solution were compared in an absorption at 450 nm. (C) SEM images of Col-I, Col-I+PEG, Col-I+PEG+HA. Scale bar measures 5 µm.

**Figure 5:**
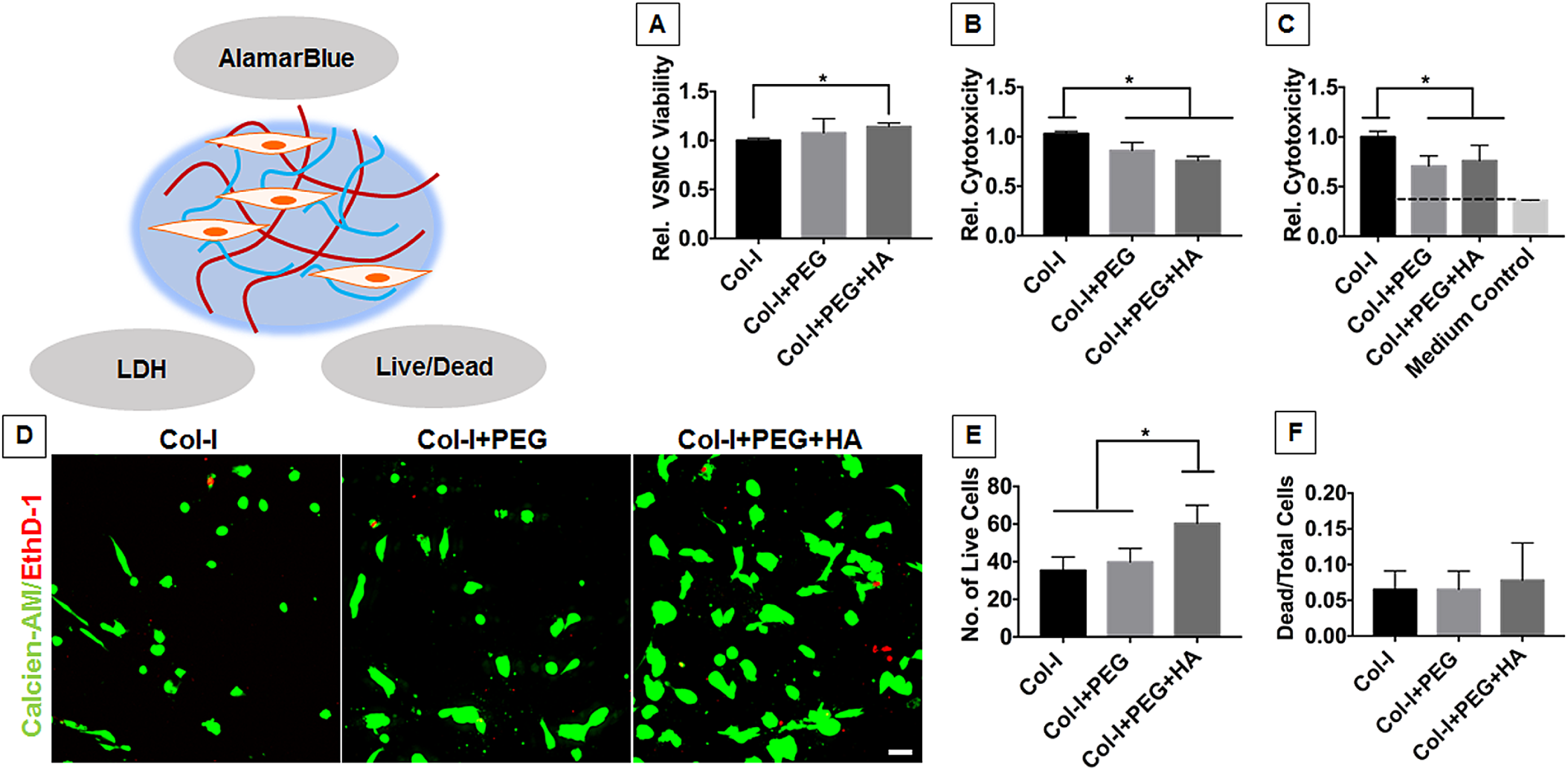
Characterization of hiPSC-VSMC viability embedded in the in situ hydrogel. (A) AlamarBlue cell viability assay showing relative cell viability of hiPSC-VSMCs embedded in the in situ Collagen-HA hydrogels of 4mg/ml of collagen concentration, 1:1 ratio of collagen to PEG cross-linker and HA of 1mg/ml final concentration on day 3. LDH assay of (B) cell lysate and (C) conditioned medium of the in situ Collagen-HA hydrogel. (D) Live/Dead assay of the in situ Collagen-HA hydrogel. Calcein-AM and EthD-1 stains live cells (Green) and dead cells (Red) respectively. Scale bar measures 50 µm. Collagen hydrogels with and without cross-linking were kept as controls for all the experiments. * denotes statistical significance differences between the different groups (n=3-6, one-way ANOVA, *p<0.05).

Furthermore, brightfield and immunofluorescence images were captured to characterize the morphology and phenotype of hiPSC-VSMCs in the hydrogels (Figure 6). Both brightfield and fluorescence images showed an elongated morphology of hiPSC-VSMCs inside Col-I+PEG and Col-I+PEG+HA hydrogels compared to uncross-linked Col-I hydrogels (Figure 6A-C). The hiPSC-VSMCs maintained their smooth muscle cell phenotype in the hydrogels as shown by staining with calponin and SM-22α (Figure 6B and C) during 3 days in vitro culture. Moreover, during this time the in situ hydrogels resisted hiPSC-VSMC mediated contraction.

**Figure 6:**
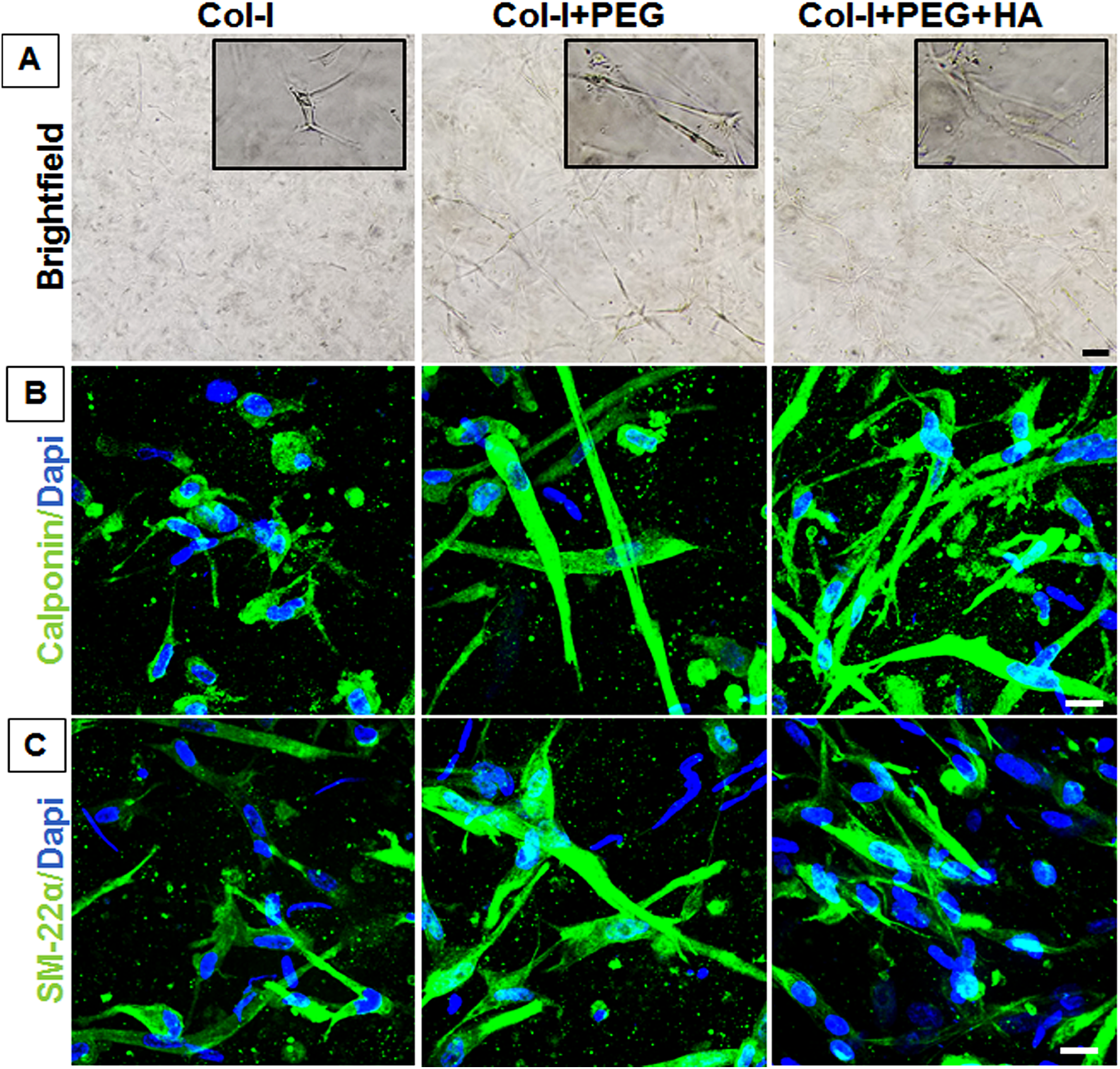
Phenotype assessment of hiPSC-VSMCs embedded in the in situ hydrogels. (A) Brightfield images of hiPSC-VSMCs embedded in the Col-I, Col-I+PEG, Col-I+PEG+HA hydrogels. Inset shows enlarged images of hiPSC-VSMCs. Scale bar measures 100 µm. Immunofluorescence images showing (B) Calponin (Green) and (C) SM-22α (Green) stained hiPSC-VSMCs. Dapi (Blue) was used to stain nuclei. Scale bar measures 20 µm.

Qualitative ELISA-based analysis demonstrated the relative level of proangiogenic growth factors VEGF, bFGF, SDF-1α and PDGF-AA in the CM (Figure 7A-D). There was no significant difference in the relative level of growth factors between any of the groups. Figure S2A shows no difference in the relative level of VEGF expression between control and various crosslinking ratios and also no difference was found between control Col-I and Col-I+HA hydrogels (Figure S2B). Functional angiogenic assay of the CM was carried out to determine the bioactivity of the secreted paracrine growth factors from the final Col-I+PEG+HA hydrogels (Figure 8) and was compared with SmGM-2, positive control endothelial cell growth medium-2 (EGM-2), and negative control endothelial basal medium (EBM). Cell adherence of HUVECs was found to be significantly more in CM compared to control SmGM-2 and EBM medium controls. The amount of adherence was however found to be at a similar level with the EGM-2 medium (Figure 8A and B). In addition, the HUVECs showed typical cobblestone-like morphology in the case of both CM and EGM-2 after 3 h of seeding. A similar pattern can be seen in the case of migration of HUVECs towards CM in a transwell migration assay (Figure 8C and D). HUVECs in the case of SmGM-2 and EBM showed reduced migration compared to CM and EGM-2 (Figure 8C and D). In vitro angiogenesis assay in Matrigel showed higher nodes/field in the case of both EGM-2 and CM and were significantly more than that of the SmGM-2 and EBM (Figure 8E and F).

**Figure 7:**
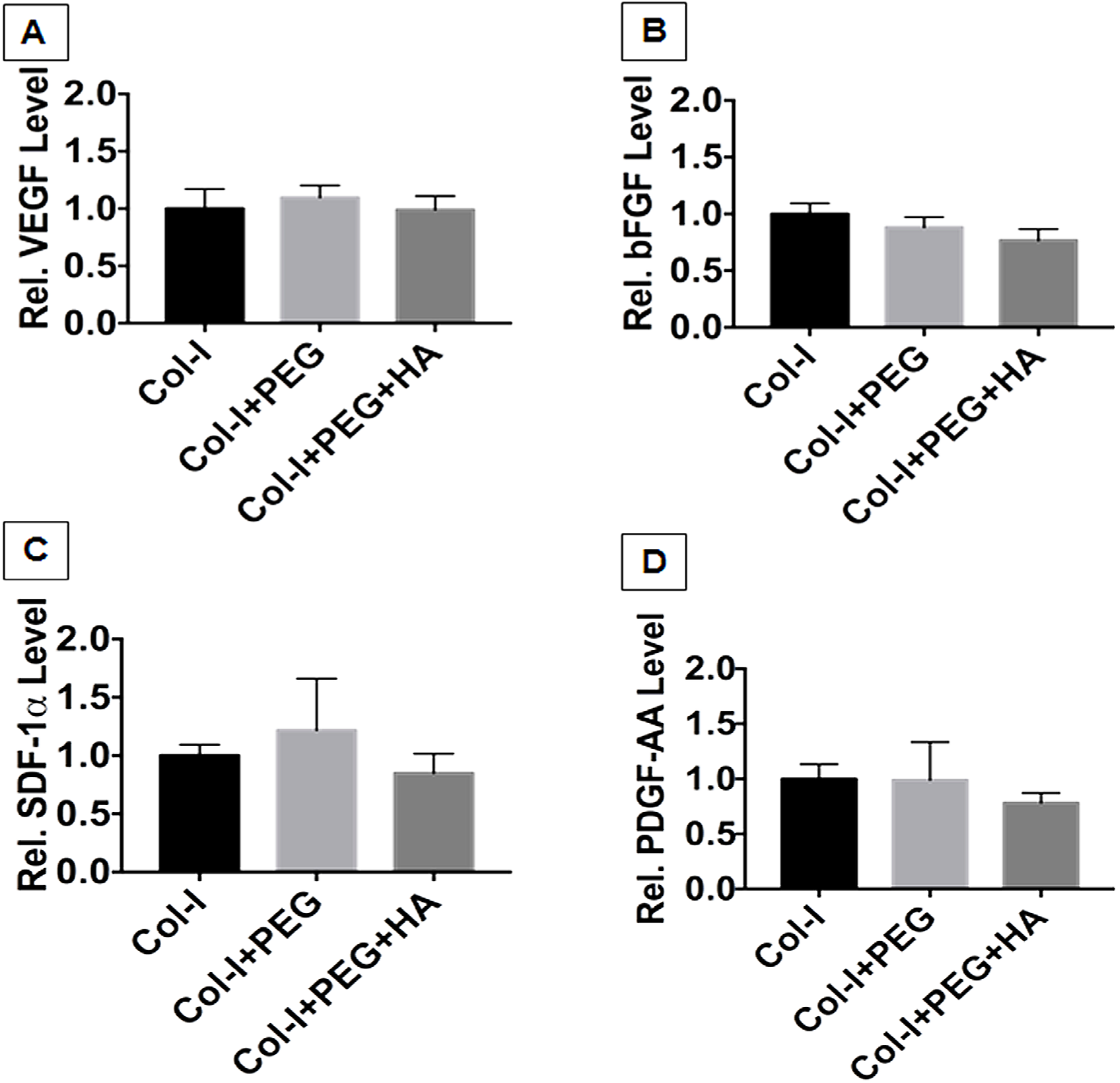
Characterization of pro-angiogenic growth factor secretion from the in situ hydrogels. Qualitative ELISA showing secretion of relative levels of growth factors (A) VEGF, (B) bFGF, (C) SDF-1α and (D) PDGF-AA in the condition medium. Collagen hydrogels with and without cross-linking were kept as controls.

**Figure 8:**
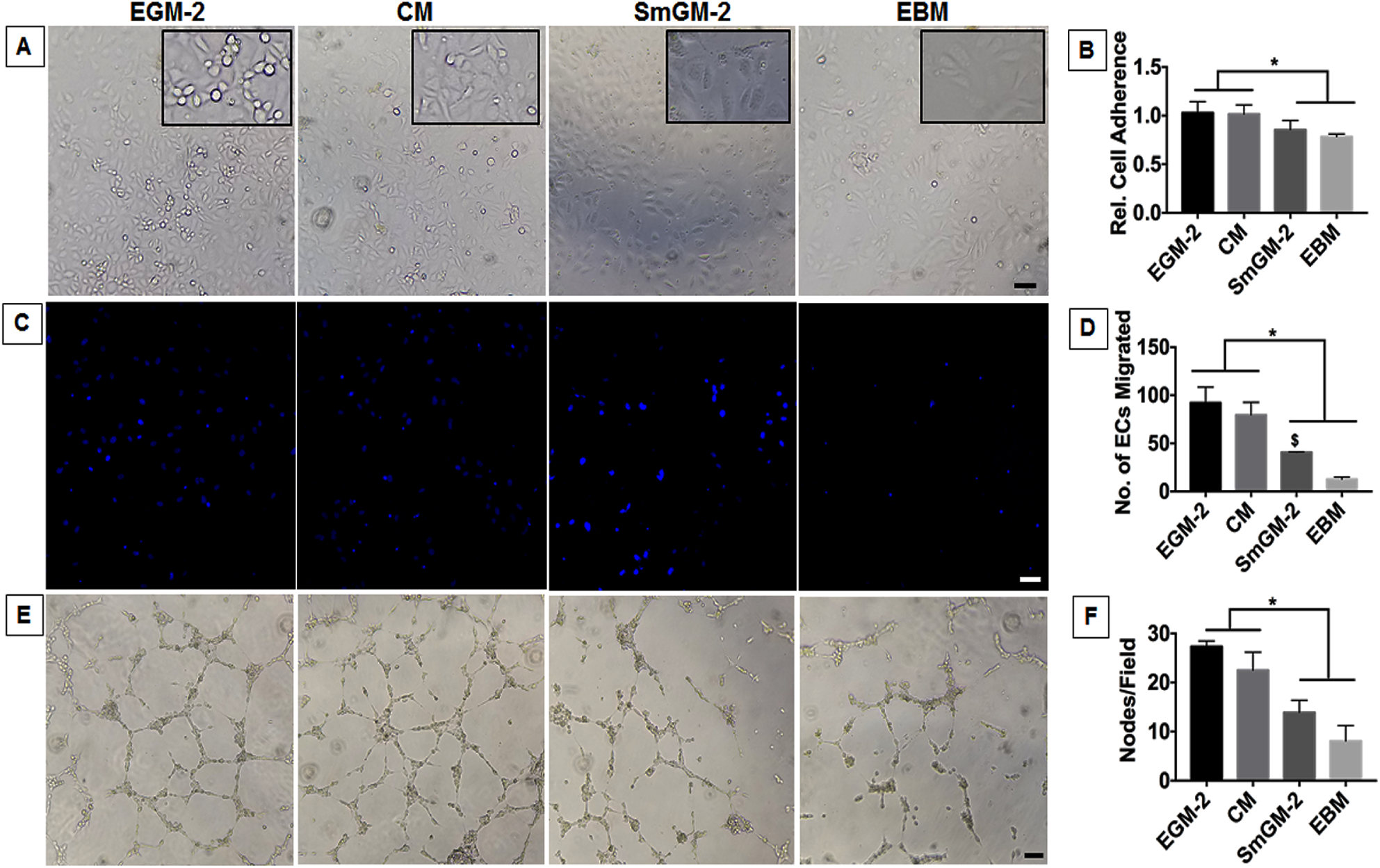
Assessment of the conditioned medium from Collagen-HA in situ hydrogels for angiogenic potentials. The conditioned medium (CM) collected from the collagen-HA hydrogels were tested for their proangiogenic bioactivity using HUVECs. (A) Brightfield images showing attachment of HUVECs on 0.1% coated gelatin wells after 3 h of seeding. Inset shows enlarged images of the HUVECs. Scale bar measures 100 µm. (B) AlamarBlue assay was performed to determine relative level of HUVECs adherence on gelatin coated plate after 3 h of seeding. Eight micrometer pore size trans-wells were used to determine migration of HUVECs in response to the CM over 4 h of incubation time. (C) The cells migrated were stained with dapi and (D) counted to obtain the total number of cells migrated. Scale bar measures 50 µm. Angiogenesis assay was performed using Matrigel and HUVECs. (**E**) Brightfield images showing tube formation of HUVECs in response to CM from in situ Collagen-HA hydrogels after 6 h of incubation. Scale bar measures 100 µm. (F) The graph represents number of nodes/field. Endothelial cell (EGM-2) and smooth muscle cell (SmGM-2) growth medium, and endothelial cell basal medium (EBM) were used as controls. * denotes statistical significance differences between the different groups (n=3, student t-test, *p<0.05).

## Discussion

Vascular smooth muscle cells derived from integration-free human iPSCs are transforming the field of vascular tissue engineering and regenerative healing with enormous translational potential (B. C. Dash, Z. Jiang, C. Suh, & Y. Qyang, 2015a; Gorecka et al., 2019). Most recently, our studies revealed the proangiogenic and anti-inflammatory potential of these cells and their use in the treatment of acute and chronic wounds (Dash et al., 2020; Gorecka et al., 2020). While these reports further confirm the future translational implications of hiPSC-VSMCs, we see an unmet need for a suitable delivery system for hiPSC-VSMCs. Our previous findings extensively studied the applicability of plastic compressed collagen scaffolds as a topical delivery system (Dash et al., 2020; Gorecka et al., 2020). However, the degradation, mechanical strength, and ease of delivery are key concerns. Here, we attempted to develop collagen and HA-based in situ hydrogels that can alleviate these issues. Furthermore, these in situ hydrogels would be appealing for wound healing applications because of their non-invasive nature. In addition, they can be considered as potent bioadhesive materials, sealant, or hemostatic agents while facilitating rapid healing.

In this study, we developed an in situ hydrogel using a combination of rat tail collagen type-I and HA for the delivery of hiPSC-VSMCs. To increase the stability of the hydrogel, collagen was cross-linked with 4S-StarPEG without compromising cell viability. The 4S-StarPEG with four NHS groups can cross-link with four amine groups on type-I collagen at 37°C and promote in situ gel formation. We determined the effectiveness of this in situ crosslinking via TNBSA quantification, which indicated a decrease in free amine groups with an increase in collagen to the 4S-StarPEG ratio. However, the effectiveness and efficiency of 4S-StarPEG cross-linking is collagen concentration-dependent. The highest degree of cross-linking was achieved in 4mg/ml of collagen hydrogels and was around 70% compared to 2.5mg/ml (∼50%) and 1.25mg/ml (∼20%). While an inefficient cross-linking found in the case of 1.25mg/ml can be attributed to less availability of amine molecules, there was no difference in cross-linking between 1:1 and 1:2 in the case of 4mg/ml hydrogels, which can be attributed to a steric hindrance caused by a saturation of 4S-StarPEG. Previous studies with both collagen I and II have demonstrated an increase in mechanical strength, viscosity, and resistance to collagenase degradation (Collin et al., 2011; Lotz et al., 2017). One such study demonstrated the use of a 4S-StarPEG cross-linked collagen scaffold as a skin substitute (Lotz et al., 2017). The report showed the development of a mechanically stronger collagen scaffold with increased resistance to collagenase degradation. Subsequent functionalization of this collagen hydrogel with HA made it more skin mimetic with regards to ECM content and strength. However, the HA was merely entrapped within the Col-I+PEG hydrogel and not cross-linked by 4S-StarPEG. The cross-linker visibly increased the transparency of in situ hydrogels as reported earlier in the case of collagen (Lotz et al., 2017). The transparent hydrogel system would help better visualization of cell morphology and wound regeneration.

After the initial cross-linking characterization of the collagen hydrogels, the cellular response to the cross-linking was assessed. We used two different primary cell types, HUVECs and dermal fibroblasts from a human source, and plated them on the top of the hydrogels to assess the viability. We saw an increase in cell viability over time for both cell types. The amount of cross-linking did not have much impact on the viability of either cell type. However, collagen concentration had a greater impact on cell viability. This is consistent with our earlier report and can be attributed to the collagen fibrillar density (CFD) (Dash et al., 2020). The CFD increases with increasing concentration of collagen and impacts cell viability and proliferation. In addition, we observed a greater amount of proliferation in the case of HUVECs compared to dermal fibroblasts. This along with the reduced proliferation of fibroblasts in the cross-linked 4mg/ml hydrogels suggest that HUVECs and dermal fibroblasts respond differently to biomechanical cues. Overall, higher concentrations of 4S-StarPEG did not compromise cell viability of primary cell types such as HUVECs and human dermal fibroblasts.

Next, we determined the effect of 4S-StarPEG on the hiPSC-VSMCs when the cross-linking reaction is conducted with the presence of embedded hiPSC-VSMCs. Similar to HUVECs and dermal fibroblasts, embedded hiPSC-VSMCs showed an increase in cell viability with increasing concentration of collagen. The viability and proliferation of embedded hiPSC-VSMCs were not affected by the cross-linker unlike earlier reports (Lotz et al., 2017). Instead, in the case of 2.5mg/ml, an increase in cell viability can be seen with an increase in amount of cross-linker. These data indicate the importance of cell-matrix interactions in cell viability and proliferation, and suggest that CFD influences viability. We believe at a lower concentration, the cell-binding motifs are less available while at 2.5mg/ml their availability is enhanced due to increased cross-linking. With regards to 4mg/ml, there are more than enough binding motifs and hence the oversaturation with cross-linking did not have an effect. This is further suggested by the observations after addition of HA to the system. Most possibly HA brings a new cellular binding motif and further increases the viscosity irrespective of the cross-linking. That may explain why an increase in cell proliferation across the various concentrations of cross-linked collagen can be seen with the addition of HA. However, the stiff increase in proliferation in the case of 1.25mg/ml can be attributed to both an increase in viscosity and additional cellular motifs.

After careful evaluation of the effects of 4S-StarPEG and HA with various combinations of collagen concentration, we chose 4mg/ml of collagen scaffold with 1:1 collagen to 4S-StarPEG cross-linker ratio and HA for the generation of robust in situ collagen/HA hydrogels. This combination not only has a higher amount of cross-linking to provide a robust skin equivalent but was associated with significant cell viability for all the cell lines tested including hiPSC-VSMCs. We saw a further increase in cell proliferation with reduced cytotoxicity in the in situ hydrogels of 4mg/ml with the addition of HA.

The hiPSC-VSMCs across all the hydrogels maintained their phenotype, however, the change in the morphology to a much-elongated shape in the cross-linked hydrogels can be due to an increase in stiffness. Furthermore, unlike the scaffolds with a lower concentration of collagen, these scaffolds do not contract when cultured with hiPSC-VSMCs over the culture period.

Although the capability of hiPSC-VSMCs to promote angiogenesis in a collagen scaffold has been illustrated (Dash et al., 2020; Gorecka et al., 2020), the effect of 4S-StarPEG cross-linking and functionalization with HA is not known. In addition, most of the cell delivery or cell-responsive platforms investigated the ability of the scaffolds to maintain cell viability and not their effect on their ability to secrete paracrine factors. This study investigated both viability and the proangiogenic response of cells in the in situ scaffolds. Here, we investigated whether cross-linking and HA will affect the secretion. Our data suggested cross-linking did not promote or adversely affected the secretion of VEGF, bFGF, SDF-1α, and PDGF-AA. This might be due to a cross-linker mediated disruption in the collagen fibril formation as shown in the SEM images. This phenomenon has already been described in published studies (Lotz et al., 2017; Yunoki & Matsuda, 2008). Similarly, HA showed no effect on paracrine secretion. We believe that the high molecular weight and negatively charged HA might be sequestering the released growth factors inside the in situ collagen-HA hydrogel. This capturing and/or depletion of cytokines and growth factors of CM through their binding to glycosaminoglycans such as HA, heparan sulfate and heparin containing ECM networks is well-described in several studies (Friedemann et al., 2017; Prokoph et al., 2012). Furthermore, we characterized the bioactivity of the secreted growth factors. Our extensive functional angiogenesis study shows an enhanced EC adherence, migration, and tube formation in response to CM. We have now shown that maintenance of growth factor production is critical as cell-mediated regenerative healing is mediated, in part, by cellular paracrine functions. This experimental inquiry demonstrated the usefulness of this delivery system as protection not only at the cell viability level and but also at the secretory level. Together, these results support the utility of a cell-laden in situ hydrogel that could be used for regenerative wound healing without any adverse effect on cell viability or production of angiogenic factors, suggesting such cellular constructs may be exploited to deliver hiPSC-VSMCs locally for wound repair applications.

## Conclusion

In summary, our initial characterization showed that 4S-StarPEG cross-linked in situ collagen type I-HA hydrogel promoted cell viability and maintained the phenotype and proangiogenic paracrine secretion of hiPSC-VSMCs. In addition, the CM from the in situ hydrogels were shown to be bioactive by promoting angiogenesis. A potential correlation between collagen density and cross-linking and cell viability can be drawn and is key to maintaining the cell viability in addition to the presence of HA. Future in-depth studies should be performed to characterize this hydrogel as a promising candidate as a carrier of hiPSC-VSMCs for wound healing applications in animal models.

## Supplementary Information

The Supporting Information is available on online and contains additional information on materials, immunostaining, optical characterization, cell culture, LDH assay, LiVE/Dead assay, In vitro angiogenesis assay, cell adherence, migration assay, alamarBlue, hydrogel fabrication, TNBSA, and ELISA. Data can be made available upon request.

## Author Contributions

B.C.D. conceived the study. B.C.D. and H.C.H., procured the funding. B.C.D. and K.D. designed the experiments. K.D., B.C.D. and H.X. performed the experiments. B.C.D. and K.D. wrote the manuscript. All the authors participated in data analysis, discussed the results and reviewed the manuscript.

## Acknowledgement

The authors would like to thank the core research facilities at Yale Department of Surgery.

---

**Schematic 1:**
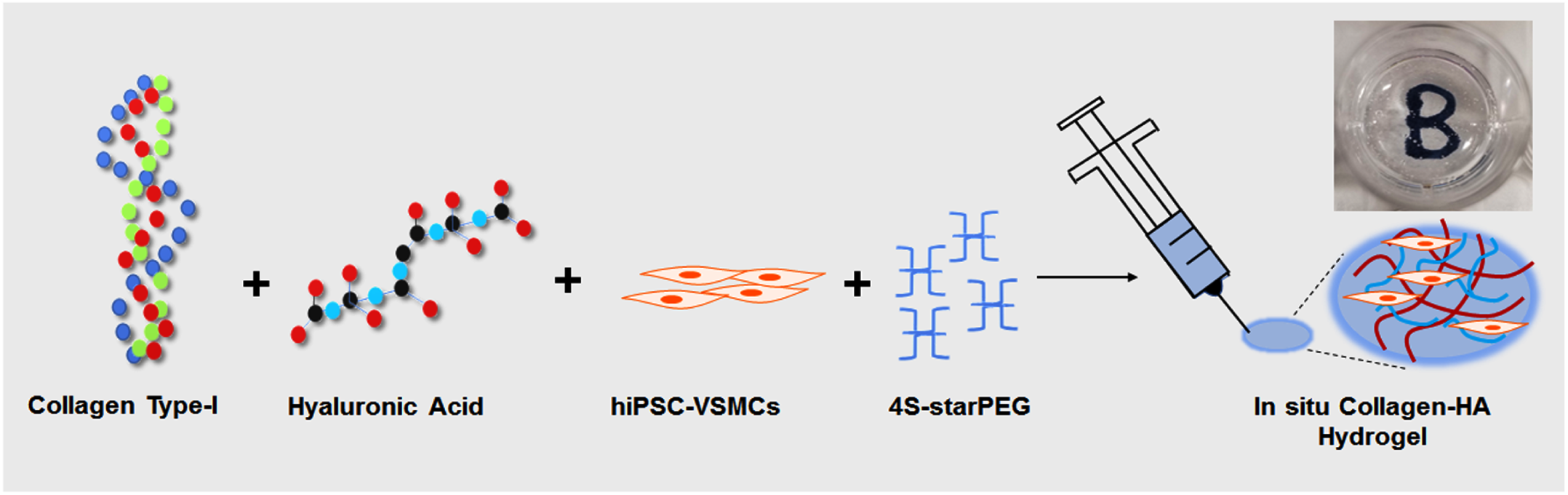
Schematic showing the fabrication of an in situ Collagen/HA hydrogel for the delivery of hiPSC-VSMCs.

