## Supplemental Information for "An In situ Collagen-HA Hydrogel System Promotes Survival and Preserves the Proangiogenic Secretion of hiPSC-derived Vascular Smooth Muscle Cells"

Running Title: In situ collagen-HA hydrogel for hiPSC-VSMCs

**^#^** Authors contributed equally

Address: Yale University School of Medicine, 330 Cedar St., BB 3^rd^ Floor, PO Box 208041, New Haven, Connecticut 06510, Telephone: 203 785-2571, Fax: 203 785-5714

**Materials:**

Type I rat tail collagen type-I was purchased from Enzo Life Sciences, USA. High molecular weight sodium hyaluronate (HA) was purchased from ACROS Organics^TM^, Fisher Scientific, USA. 4-arm polyethylene glycol succinimidyl glutarate Mw 10,000 (4S-StarPEG) was purchased from JenKem Technology, USA. Trinitrobenzene sulfonic acid (TNBSA) and AlamarBlue reagents were purchased from ThermoFisher Scientific, USA. Smooth muscle cells culture medium (SmGM-2) was purchased from PromoCell, Germany. All other cell culture reagents were purchased from ThermoFisher Scientific unless otherwise stated.

**2,4,6-trinitrobenzene sulfonic acid assay:**

2,4,6-Trinitrobenzene Sulfonic Acid (TNBSA) assay was performed to determine the degree of crosslinking of collagen type I as per the previously method (Collin et al., 2011). The efficacy of 4S-StarPEG of crosslinking collagen type-I was tested in varies collagen densities (1.25mg/ml, 2.5mg/ml, and 4mg/ml). The degree of PEG cross-linking in varying Collagen to PEG ratios (control, 1:0.5, 1:1, and 1:2) was also evaluated. Glycine titration in sodium bicarbonate (0.1M) pH 8.5 standard curve was established at: 300uM, 250uM, 200uM, 150uM, 100uM, 75uM, 50uM, 25uM, and 0uM. 250uL of 0.01% TNBSA was added into 500uL of each sample. A 0.2% glutaraldehyde solution was used as a positive control. The resultant mixtures were incubated at 37^o^C for 2 hours. After incubation, 250µL of 10% sodium dodecyl sulfate was added. Then, 125µL of 1M HCI was added to and mixed to stop the reaction. 100µL of the final solution was used to test for the degree of amine group cross-linkage. Plate reader absorbance was set at 335nm. Percentage of cross-linking was evaluated by considering the amount of free primary amine groups in control collagen solution as 100%.

**Optical density characterization of in situ hydrogels:**

For optical imaging uncross-linked collagen hydrogels and cross-linked collagen hydrogels with and without HA were plated in a 24 well plate. Photos were captured using a digital camera. The optical density of hydrogels of 100 μl of the volume was determined by absorption of 450 nm in a 96-well plate using a Tecan Infinite M200 plate reader. Percent transmittance was calculated using the following formula: % Transmittance = antilog (2 – absorbance).

**Scanning Electron Microscopy:**

For scanning electron microscope experiments Col-I, Col-I+PEG, and Col-I+PEG+HA hydrogels were first fixed in 2.5% paraformaldehyde in 0.1M cacodylate buffer for 15 min. This step was followed by dehydration of the samples using an ethanol gradient of 50%, 60%, 70%, 80%, 90%, 100%, 100% for 5 min each. The dehydrated samples were then incubated in hexamethyldisilazane (Electron Microscopy Sciences) and air-dry overnight. Samples were mounted onto the sample stage using carbon tapes and then coated with 8 nm Iridium using a Cressington 208 iridium sputtering tool. SEM images were taken on a Hitachi SU-70 scanning electron microscope using a 5 kV acceleration voltage.

**Cell culture:**

Human primary cells: Human umbilical vascular endothelial cells (HUVECs) and human dermal fibroblasts of passage number below 7 were used in the experiments. Both the primary cells were cultured on 1% gelatin-coated plates. The medium for HUVECs was endothelial cell growth medium (EGM)-2 and for fibroblasts, it was DMEM high Glucose with 10% FBS. The respective medium was replenished with a fresh one every other day and cultured until 80% confluency before using them on hydrogels.

hiPSC-VSMCs: hiPSC-VSMCs were differentiated using a previously described protocol (Dash et al., 2016). Briefly, hiPSCs were initially cultured under feeder-free conditions until reaching 80% confluency in a 6-well culture plate (4 days). Then clumps of uniform sizes were harvested after treating with dispase for 15 minutes at 37^o^C. The resulting hiPSC clumps were then seeded in a 6-well low attachment plate with mTESR medium. Next day, the medium was replaced with mixture of mTeSR and EB differentiation medium (DMEM high glucose + 10% FBS + 1% non-essential amino acid (v/v) + 2mM L-glutamine and 0.012 mM 2-mercaptoethanol) at a 1 to 3 ratio. On day 3, the medium was replaced with just the EB differentiation medium. On day 5, the resulting EB were collected and seeded on a gelatin-coated 6-well plate and cultured in EB differentiation medium for 5 days or until 80% confluency (around 7 days). After reaching the targeted density, the smooth muscle cell growth medium (SmGM)-2 medium was used for culturing and was replaced every other day. The resulting differentiated cells were validated with immunofluorescence staining with calponin and SM-22α. Human iPSC-VSMCs were cultured until at least 80% confluence, then they were seeded into collagen-based hydrogels for subsequent experiments.

**Immunofluorescence staining:**

Human iPSC-VSMCs cultured on tissue culture plates and scaffolds were stained with anti-calponin and anti-SM-22α primary antibodies for phenotype characterization. First, the samples were blocked with 5% BSA in 0.25% Triton X for 1 hour at room temperature. Subsequent primary antibodies were added for incubation overnight at 4^o^C. On the next day, the scaffolds were washed three times with PBST (tween20 0.05%), and then the scaffolds were incubated with secondary antibodies tagged with Alexa Fluor® 488. Dapi was used as a counterstain. Z stacks were taken at the interval 1 µm and images were merged together for the final image. Lastly, the samples were washed with PBST three times at 5 minutes each before fluorescence imaging. The information related to primary and secondary antibodies can be found in table S4 and S5.

**AlamarBlue assay:**

The cellular collagen scaffolds containing hiPSC-VSMCs were characterized for cell viability on day 3. HUVECs and dermal fibroblasts cultured on top of hydrogels were evaluated for viability on days 1 and 3. Briefly, 200µl of culturing media was gently aspirated out for future ELISA assays. The scaffolds were then gently washed with 200µl of PBS. AlamarBlue stock was made at a 1:10 ratio of alamarBlue to medium. 100µl of the stock solution was added to each of the hydrogels and incubated at 37^o^C for 2 hours. At the end of the incubation, the plate was read for Fluorescence intensity at 540nm excitation and 590 emission wavelengths. Relative cell viability was evaluated by dividing them with the collagen scaffold fluorescence intensity value.

**Lactate dehydrogenase assay:**

Lactate dehydrogenase cytotoxicity assay was performed using an LDH cytotoxicity assay kit (Peirce, Thermofisher) as described (Dash et al., 2020). Briefly, conditioned medium (CM) and cell lysates were used to determine the level of LDH in different samples. For lysates, the hydrogels were homogenized using mechanical force and RIPA buffer. The cell lysates and CM were then used to detect LDH levels as per the manufacturer’s instruction. The different groups that were included in this experiment were: Col-I, Col-I+PEG, and Col-I+PEG+HA. Absorbance was measured at 490 and 680nm. To determine LDH activity, the 680nm absorbance value (background signal from the instrument) was subtracted from the 490nm absorbance. Relative cytotoxicity value to control Col-I was determined.

**Live/Dead assay:**

Cell viability was assessed using the Live/Dead assay. Briefly, hydrogels were incubated in PBS supplemented with 4 μM calcein-AM green (ThermoFisher, USA) and 2 μM ethidium homodimer-1 (ThermoFisher, USA) for 30 min. Stained samples were visualized on an inverted Confocal Laser Scanning Microscope. Z stacks were performed at 1 μm image intervals and images were merged for the final image. Four fields per hydrogel, with three hydrogels per condition, were imaged for each experiment. Viable and dead cells were counted using ImageJ software. The number of live and dead cells was then estimated.

**Qualitative ELISA:**

Day 3 CM that were collected from the hydrogels were used to perform qualitative ELISA to evaluate for several pro-angiogenic growth factors VEGF, bFGF, SDF-1α, and PDGF-AA released from hiPSC-VSMCs as described (Dash et al., 2020). 90µl of the CM was used to microwells of 96 well ELISA plates (NUNC MaxiSorp™). The plates were incubated at 4^o^C overnight. The next day, the plates were washed with PBST (PBS + 0.05% Tween-20) three times and were blocked with 5% BBSA for 1 hour at 37^o^C. Then the plates were washed with PBST three times and were incubated with primary antibodies (1:2500) at 4^o^C overnight. The plates were washed three times with PBST and incubated with secondary antibodies conjugated with horseradish peroxidase (1:2500) for 2 hours at room temperature. The plates were then washed 3 times with PBST, 100µl of TMB substrate solution (Cell Signaling Technology, USA, Catalog: 7004S) added to the plates after aspirating out the PBST and was incubated for 25 minutes at room temperature on a plate shaker. 100µl of stop solution (Cell Signaling Technology) was then added to each microwell and absorbance was measured at 450nm on a plate reader. The information related to primary and secondary antibodies can be found in table S4 and S5 in the supplementary information. Relative level of growth factor was evaluated by dividing them with the collagen scaffold absorbance value.

**Cell adherence and migration assay:**

Human umbilical vein endothelial cells (HUVECs) were grown until 80% confluency. The cells were then dissociated using TrypLE and were used for subcultures and cell adherence and migration assays. Cell adherence assays for HUVECs was performed in a 96 well plate. Briefly, 10,000 cells were plated on a 96 well plate and cultured for 3 h in CM supplemented endothelial basal medium (EBM) of a ratio of 1:5. Controls were endothelial growth medium-2 (EGM-2), smooth muscle cell growth medium (SmGM-2), and EBM. For migration, transwell assays were performed using HUVECs as described (Dash et al., 2020). Briefly, 1 x 10^4^ cells were seeded on the top layer of the trans-wells (8µm pore size trans-wells, Falcon) and incubated for 10 min in cell culture incubators. The trans-wells were then placed in 24 well plates and 600µl of respective growth medium with CM were added to the wells touching the inner side of the trans-well. The trans-wells were then kept in the cell culture incubator for 4 h. This was followed by fixing with 100% cold methanol and gently removing the cells from the top layer. Trans-wells were washed and stained using dapi. The number of cells migrated to the inner side of the membrane was then quantified by counting the number of dapi stained nuclei from 4 different fields of view.

**In vitro angiogenesis assay:**

The ability of hiPSC-VSMC-CM from in situ hydrogels to induce angiogenesis was measured using an in vitro angiogenesis assay as described (Dash et al., 2020; DeCicco-Skinner et al., 2014). HUVECs were cultured until they were 80% confluent in EGM-2 medium. A 100 µl/well of Matrigel (Corning Life Sciences) was spread evenly over the wells of a 96-wells plate. The plates were incubated for 30 min at 37℃ to allow the Matrigel to solidify. HUVECs (2x10^5^/ml) were resuspended in EGM-2 basal medium and 100µl of this along with 100µl of CM were seeded in each well (n=3). After 6 h of incubation of the plate at 37^o^C, the floating cells were removed & the plates were washed (2x) with PBS and fixed using PFA for brightfield imaging. Images from 4 different fields/well were acquired and the number of nodes per field was counted.

**Table S1:** Collagen-4S-StarPEG hydrogel preparation for cells on the top.

| Density (mg/ml) | Collagen (μl) | 10x MEM  (μl) | 1M NaOH  (μl) | 1x PBS  (μl) | 4S-StarPEG 25nM  (1:0.5 ratio)  4S-StarPEG | 4S-StarPEG 25nM (1:1ratio)  (μl) | 4S-StarPEG 25nM  (1:2 ratio)  (μl) |
| --- | --- | --- | --- | --- | --- | --- | --- |
| 1.25 | 125 | 50 | 2.3 | 325 | 2 | 4 | 8 |
| 2.5 | 250 | 50 | 3 | 200 | 4 | 8 | 16 |
| 4 | 400 | 50 | 9 | 50 | 8 | 16 | 32 |

**Table S2:** Collagen-4S-StarPEG hydrogel preparation for cell embedding.

| Density  (mg/ml) | Collagen  (μl) | 10xMEM  (μl) | 1M NaOH  (μl) | SmGM-2  (μl) | VSMCs  (8000/μl) | 4S-StarPEG 25nM  (1:0.5 ratio) (μl) | 4S-StarPEG 25nM  (1:1 ratio) (μl) | 4S-StarPEG 25nM  (1:2 ratio) (μl) |
| --- | --- | --- | --- | --- | --- | --- | --- | --- |
| 1.25 | 125 | 50 | 2.3 | 300 | 25 | 2 | 4 | 8 |
| 2.5 | 250 | 50 | 3 | 175 | 25 | 4 | 8 | 16 |
| 4 | 400 | 50 | 9.0 | 25 | 25 | 8 | 16 | 32 |

**Table S3:** Collagen- 4S-StarPEG-Hyaluronic Acid in situ hydrogel preparation.

| Density  (mg/ml) | Collagen  (μl) | 10xMEM  (μl) | 1M NaOH  (μl) | SmGM-2  (μl) | VSMCs  (8000/ μl) | HA (1mg/ml)  (μl) | 4S-StarPEG 25nM  (1:0.5 ratio)  (μl) | 4S-StarPEG 25nM  (1:1 ratio)  (μl) | 4S-StarPEG 25nM  (1:2 ratio)  (μl) |
| --- | --- | --- | --- | --- | --- | --- | --- | --- | --- |
| 1.25 | 125 | 50 | 2.3 | 300 | 25 | 25 | 2 | 4 | 8 |
| 2.5 | 250 | 50 | 3 | 175 | 25 | 25 | 4 | 8 | 16 |
| 4 | 400 | 50 | 9 | 0 | 25 | 25 | 8 | 16 | 32 |

**Table-S4:** Information related to primary antibodies.

| Primary Antibody | Dilution | Catalog Number/ Manufacturer |
| --- | --- | --- |
| SDF-1α | 1:2500 ELISA | MAB350 (R&D) |
| PDGFAA | 1:2500 ELISA | 500-P46-100 (PeproTech) |
| bFGF | 1:2500 ELISA | 500M38 (PeproTech) |
| VEGF | 1:2500 ELISA; 1:200 IHC | AB-119 (Abcam) |
| SM-22α | 1:300 IF and IHC  1µg/ml FACS | AB-10135 (Abcam) |
| Calponin | 1:200 IF and IHC  1µg/ml FACS | C-2687 (Sigma) |

| Secondary Antibody | Dilution | Catalog Number/ Manufacturer |
| --- | --- | --- |
| Anti-Mouse-HRP | 1:2500 ELISA | AB6789 (Abcam) |
| Anti-Rabbit-HRP | 1:2500 ELISA | A0545 (Sigma) |
| Anti-Mouse-Alexafluor488 | 1:400 IF and IHC | A11029 (ThermoFisher) |
| Anti-Goat-Alexafluor488 | 1:400 IF and IHC | A27012 (ThermoFisher) |

**Table-S5:** Information related to secondary antibodies.

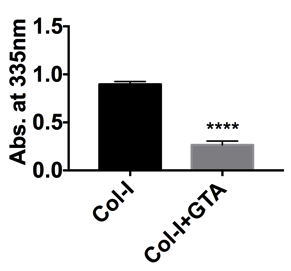

**Figure S1: Characterization of glutaraldehyde cross-linking collagen type-I.** TNBSA showing a qualitative analysis of free amine groups in crosslinked collagen hydrogel of 4mg/ml of collagen concentration. Uncross-linked collagen hydrogel was kept as a control. * denotes statistical significance differences between the different groups (n=4, t-test, ****p <0.0001).

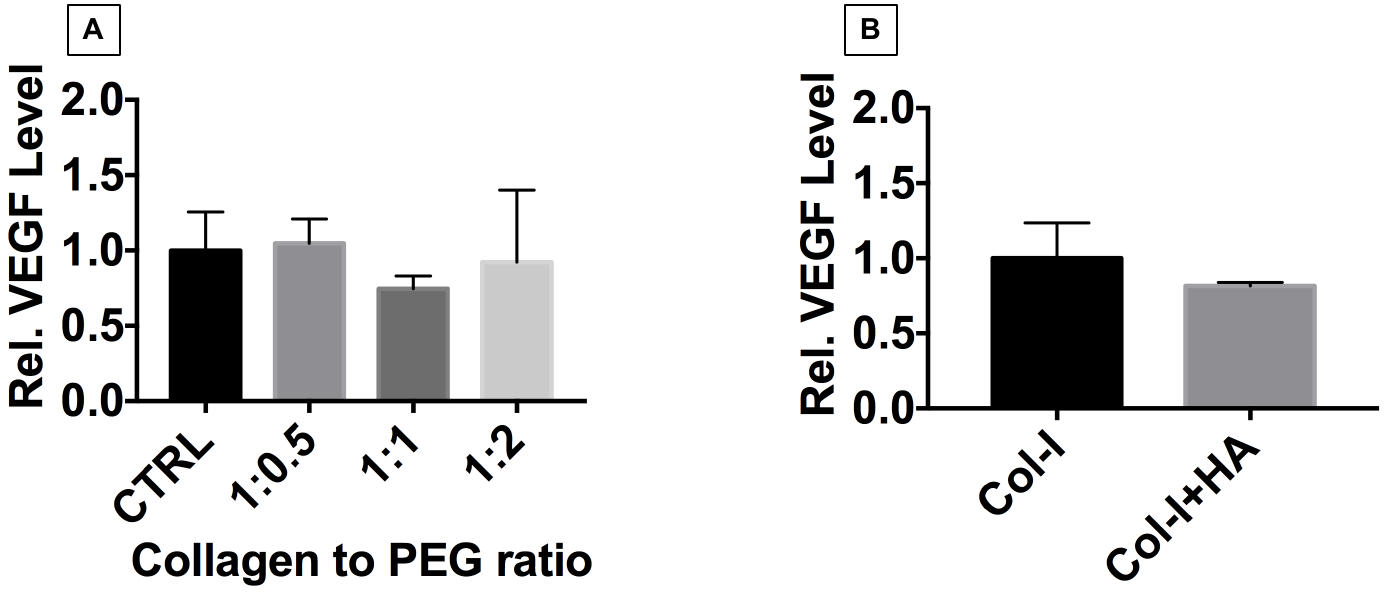

**Figure S2:** **Characterization of VEGF secretion.** Qualitative ELISA showing secretion of relative level of VEGF from (A) In situ collagen hydrogel with various molar ratios of collagen to 4S-StarPEG and (B) collagen-HA hydrogel. Uncross-linked collagen hydrogel and without HA was used as control. (n=4-6, one-way ANOVA, p>0.05).
